# Eicosapentaenoic Acid Enhances Angiogenesis and Reperfusion After Ischemia: Comparative Effects of Omega-3 Fatty Acids in a Murine Hind Limb Model

**DOI:** 10.64898/2026.02.09.704971

**Authors:** TK Dao, G Barsha, J Thomas, A Pokrassen, SJ Nicholls, KJ Bubb

## Abstract

**Background:** Omega-3 polyunsaturated fatty acids are known to confer benefits in the prevention of cardiovascular diseases. Among these, eicosapentaenoic acid (EPA) appears to be more effective than docosahexaenoic acid (DHA) in ischemic vascular conditions. However, the specific roles of EPA and DHA in limb angiogenesis and post-ischemic reperfusion remain unclear. Moreover, omega-3 fatty acids remain understudied for peripheral artery disease (PAD) intervention. The aim was to compare the effect of high dose omega-3 fatty acids, EPA and DHA on ischemic tissue reperfusion, angiogenesis and vascular remodelling in mice.

**Methods:** Hind limb ischemia (HLI) was performed in mice and reperfusion was measured using laser speckle contrast imaging over two weeks post-HLI. Mice were treated daily with oral high-dose EPA or DHA (600 mg/kg/day) or vehicle (olive oil). Gastrocnemius muscle tissue was collected for analysis of mRNA and protein markers of angiogenesis.

**Results:** Following HLI, blood flow was restored more rapidly in mice treated with EPA compared with vehicle. DHA treatment did not enhance reperfusion. Histological assessment revealed significant muscle fibre regeneration after HLI, which was further improved by EPA. CD31+ neo vessel density was also increased in the EPA group. Collectively, these findings indicate that EPA promotes angiogenesis after peripheral vascular ischemia, whereas DHA does not. The beneficial effects of EPA are associated with upregulation of hypoxia inducible factor α.

**Conclusions:** High-dose EPA accelerated post-ischemic reperfusion, while DHA was ineffective. These results highlight EPA as a potential therapeutic strategy for improving limb perfusion and vascular repair in patients with PAD.

**Research Perspective:** *What Is New?:*

- Omega-3 fatty acid, eicosapentaenoic acid (EPA) accelerates post-ischaemic reperfusion following hind limb ischemia injury in mice.
- Omega 3 fatty acid, docosahexaenoic acid (DHA), does not show the same improvement in reperfusion after hind limb ischemia
- EPA can promote vasculogenesis and stimulate muscle fibre regeneration.

*What question should be addressed next?:*

- High dose purified EPA, in the form of icosapent ethyl reduces mortality from coronary artery disease. Peripheral artery disease (PAD) is a common co-morbidity yet high quality interventional trials for icosapent ethyl for PAD are lacking. Therapeutic angiogenesis has the potential to improve PAD symptoms and disease progression but there are no efficacious candidates. Icosapent ethyl should be trialled to determine whether functional outcomes are improved in PAD.

## Introduction

Peripheral Artery Disease (PAD) is a progressive disorder that increases sharply with age, with an estimated prevalence of 10% in people aged 65 or more ^1^, increasing to 20% in ages over 80 years ^2^. The most common cardiovascular risk factors, including atherosclerosis, diabetes, smoking and hypertension contribute to restricted blood flow, often causing pain in the lower extremities. When compensatory collateral vessel development or vasodilation fails, PAD can progress to chronic limb-threatening ischemia comprising necrosis, ulceration and gangrene ^1^, and can necessitate amputation. Intervening to halt the declining perfusion could substantially improve outcomes for people with PAD. Therefore, investigational therapeutic regimes are aimed at exploiting the compensatory and complementary mechanisms of reperfusion of ischemic tissue.

Circulating levels of polyunsaturated omega-3 fatty acids have long been associated with cardiovascular disease (CVD) benefit ^3^. Omega-3 fatty acid supplements are now recommended in high doses for people with hypertriglyceridemia ^4^. Non-prescription supplements are generally formulated with similar amounts of eicosapentaenoic acid (EPA) and docosahexaenoic acid (DHA), yet it has emerged that there may be vastly greater benefit of EPA for cardiovascular health. The large-scale randomised control trial (RCT), REDUCE-IT found significantly reduced cardiovascular mortality with administration of pure high-dose EPA, ^5^ whereas, the STRENGTH RCT, which investigated a combined carboxylic acid formulation containing both EPA and DHA, was terminated early due to a low probability of reduction of cardiovascular risk ^6^. Thus, we speculated that DHA may offset the positive effects of EPA in the coronary circulation. Multiple studies spanning decades have investigated the effects of numerous formulations of omega 3 fatty acids in PAD ^7-10^. Yet the majority were low in power, and/or low in dose, and often excluded diabetic people. Therefore, it remains inconclusive as to whether high doses of omega 3 fatty acids could improve leg perfusion ^11,12^.

Post-ischemic angiogenesis is an attractive option to endogenously improve blood flow to ischemic tissue and is particularly relevant in PAD ^13^ and stroke ^14^. A well-established mechanism involves upregulated hypoxia-inducible factor alpha (HIF1α) leading to vascular endothelial growth factor (VEGF) transcription ^15^ and downstream antioxidant effects ^13,14,16^. Omega-3 fatty acids have anti-inflammatory effects and are implicated in the attenuation of oxidative stress-induced DNA damage in vascular endothelial cells ^17^. Yet their role in endothelium-driven angiogenesis is contentious. Whilst omega-3 fatty acids inhibit angiogenesis in tumours, reducing incidence of cancers ^18^, after vascular ischemia omega-3 fatty acids may promote angiogenesis to restore perfusion to downstream hypoxic tissues. For example, mice fed with a diet supplemented with EPA/DHA displayed better histological outcomes in the cerebral circulation, as compared to mice on control diets ^19^, following cerebral ischemia. Moreover, cerebral angiogenesis is increased by omega-3 fatty acids after stroke ^20^. After acute coronary syndrome (myocardial infarction), many pre-clinical studies have demonstrated attenuated cardiac remodelling, due to reduced oxidative stress and inflammation in response to omega-3 fatty acid treatment (reviewed in ^21^). Likewise, the OMEGA-REMODEL RCT demonstrated left ventricular functional improvement and reduced non-infarct fibrosis after only 1g/day of interventional mixed omega-3 fatty acid supplementation ^22^. A single study has shown improved post-ischemic reperfusion and neovascularisation in the peripheral circulation after a fish oil-enriched diet ^23^, but no studies have examined the individual roles of EPA and DHA. This significant gap in current knowledge has contributed to the sub-optimal investigation of omega-3 fatty acids for PAD and provides rationale for further pre-clinical modelling to better understand the unique effects of EPA and DHA prior to investigation in further clinical trials.

We aimed to compare the effects of high dose EPA and DHA on reperfusion and vascularisation after hind limb ischemia in mice. We hypothesised that EPA would be effective at improving post-ischemic reperfusion and angiogenesis, and that DHA would be ineffective or even inhibitory.

## Methods

### Mice and animal ethics approval

All murine studies conformed and had approval from the Monash University Animal Ethics Committee (#22704) and were conducted in accordance with the National Health and Medical Research Council of Australia Guidelines for Animal Experimentation. A total of 58 male C57BL6/J mice aged 7-12 weeks were obtained from Monash Animal Research Platform to fulfill all treatment groups with two timepoints for tissue collection (day 7 and 14). The body weight range of mice at baseline was as follows: vehicle 26.7±0.29 g (n=24); EPA 27.5±0.45 (n=19); DHA 27.7±0.57 (n=15).

### Hindlimb Ischemia (HLI)

HLI was performed as described previously ^24^. Briefly, male C57BL6 mice received pre-operative local anaesthetic, bupivacaine (4mg/kg) under the skin incision and an analgesia cocktail of buprenorphine (0.05 mg/kg, s.c) and carprofen (5 mg/kg, s.c) and were then anaesthetised with isoflurane in room air (2-5%, 400-500 ml/min). Once reflexes were absent, mice were placed in supine position over a draped sensor warming pad with rectal probe homeothermic feedback to maintain body temperature at 37°C. The breathing rate of the mouse was monitored and maintained in the range of ∼60-80 breaths/minute. A 0.5-1cm vertical skin incision from the knee towards the medial thigh was made to expose the femoral vasculature. Subcutaneous fat tissue surrounding the thigh muscle was pushed aside and forceps were used to gently pierce through the membranous femoral sheath, and the femoral artery and vein were isolated from the neurovascular bundle of femoral nerves. 6-0 sutures were placed 1) proximal and 2) distal to the origin of the profunda femoris artery and 3) proximal to the saphenous artery. The femoral artery and vein were then excised between ligatures 2 and 3. The overlying skin was closed with non-continuous 6-0 polypropylene suture (Ethicon Prolene, Johnson and Johnson Medical, Australia). Non-invasive laser speckle contrast imaging (MoorO2Flo Perfusion & Oxygenation Imager, Moor Instruments Ltd, UK) was used to assess dynamic blood flow. The limb mean flux units were quantified and expressed as a ratio of the flux value of the ischemic limb relative to the contralateral non-ischemic limb in the same mouse to control for the minor variations in body temperature and ventilation. Imaging was performed 1-3 days prior to HLI (baseline), the morning after HLI surgery (day 1) and then at days 1, 5, 7, 10 and 14 post HLI.

Mice were randomly assigned to treatment groups and were oral gavaged daily with high-dose, purified EPA or DHA (600 mg/kg/day, NuChekPrep, USA) from day 0 to 14. Control mice underwent the same HLI protocol but were treated with olive oil (vehicle used for diluting EPA and DHA). At post-mortem (Day 14), mice were briefly anaesthetised with isoflurane and blood was collected via puncture of the abdominal aorta. Mice were culled by cervical dislocation and the gastrocnemius muscle and adductor muscles for both ischemic and non-ischemic limbs were collected for later analysis.

### Hind limb histopathology

At the end of the 14-day hind limb ischemia protocol, mice were anaesthetised with isoflurane (3.5%) and euthanised by exsanguination via blood collected from the abdominal vena cava using a heparinised 23-gauge needle, followed by cervical dislocation. The gastrocnemius muscles of both the ischaemic and non-ischaemic hindlimbs were emersion fixed in formalin solution (10%) and embedded in paraffin. Gastrocnemius sections (4 µm) were stained with either Picrosirius red or hematoxylin and eosin by the Monash Histology Platform. Slides were imaged using an Aperio Slide Scanning Unit (Leica Biosystems, Wetzlar, Hesse, Germany) and images (20× magnification) were captured using Aperio ImageScope Software (Version 12.4.6.5003; Leica Biosystems, Wetzlar, Hesse, Germany; RRID:SCR_020993). Picrosirius red staining was quantified to determine percentage collagen content and muscle fibre number and centralised nuclei were determined from hematoxylin and eosin staining.

Following sodium citrate antigen retrieval (pH 6; AJAX Finechem, Taren Point, NSW, Australia), gastrocnemius sections were deparaffinised, rehydrated and probed for neo vessel (CD31+) formation similar to previously described ^25^. Briefly, sections were blocked with a solution of hydrogen peroxide (0.3% H2O2 in methanol) and goat serum (10% (v/v) goat serum in PBS) before being incubated in rabbit anti-CD31 (1:25; ab28364; Abcam, Cambridge, MA, USA; RRID:AB_726362) overnight at 4°C. Primary antibodies were detected using horseradish peroxidase (HRP) secondary antibody (1:200; ab6721; Abcam, Cambridge, MA, USA; RRID:AB_955447) and 3,3′-Diaminobenzidine (DAB) substrate (Vector Laboratories, Burlingame, CA, USA). Cell nuclei were counterstained with haematoxylin and images were scanned and imaged using the Aperio system as outlined above. Image analysis was conducted by individuals blinded to the sample treatment. For all sections multiple images (minimum of 5) were taken over each section and analysed using Image J version 1.5. Average values were calculated and plotted.

### Quantitative RT-PCR (q-PCR) of gastrocnemius muscle

RNA was extracted from gastrocnemius muscle tissue using the RNeasy® Mini Kit (Qiagen, Australia). mRNA concentration was determined using the Nanodrop^™^ One/One^C^ Microvolume UV-Vis Spectrophotometer (ThermoFisher Scientific, Australia). 1 μg of mRNA was converted to cDNA using the QuantiTect Reverse Transcription Kit (Qiagen, Australia). 20 ng of cDNA from each sample was used for quantitative reverse transcription polymerase chain reaction (RT-qPCR) using the QuantiTect® SYBR® Green PCR (Qiagen, Clayton, Australia). The following KicQStart primers (Sigma, Australia) were used (500 nM): VEGF forward TAGAGTACATCTTCAAGCCG, reverse TCTTTCTTTGGTCTGCATTC, HIF1α forward CGATGACACAGAAACTGAAG, reverse GAAGGTAAAGGAGACATTGC and β-actin (housekeeping gene) forward GATGTATGAAGGCTTTGGTC, reverse TGTGCACTTTTATTGGTCTC. PCR was amplified over 40 cycles using the QuantStudio™ 7 Real-Time PCR system (ThermoFisher Scientific, USA) and data was captured and analysed using Design and Analysis Software 2.7.0. mRNA expression was normalised to β-actin and each sample was expressed as 2-ΔΔCt relative to the average of the non-ischemic vehicle control group.

### Cell assays

Human umbilical vein endothelial cells (HUVEC) were obtained from a pooled source of donors from a commercial supplier (EGM-2; Lonza, Switzerland) and cultured in endothelial growth medium 2, containing 2% bovine serum albumin (Lonza, Switzerland) between passage 3-8. For all assays cells were re-suspended in EGM2 diluted 1:3 in basal medium to reduce the serum and growth factors to 25%, which does not affect morphology ^24^. For scratch migration assays, cells were seeded at 2×10^4^ cells/well in 96 well plates (Corning, UK) and left overnight to reach confluence. The cell monolayer was scratched using a 10 µL sterile pipette tip, and images were taken at 0h, 2h, 4h, 6h and 8h to monitor scratch closure. For tubule formation assays, HUVECs were seeded at 0.5×10^4^ cells/well in 96 well plates on a layer of basement membrane extract (reduced growth factor, Pathclear, Cultrex, R&D Systems Inc., USA). After 16 hours, the number of tubules was quantified using Wimasis platform (Onimagin Technologies, Spain). Cell populations were treated with either EPA or DHA (10 µg/ml, #90110 and #90310 Cayman Chemicals, USA) or a combination of both at the time of scratching or seeding on extracellular matrix and closure rates were compared to vehicle treatment (0.1% ethanol).

### Experimental design and Data Analysis

Mouse studies were powered to achieve probability of detecting a meaningful effect at 0.8 with alpha significance level of 5% (p<0.05), with an effect size of 0.15 considered biologically relevant for the primary endpoint of flux ratio between ischemic and non-ischemic limb. Based on standard deviation of previous experiments of 0.1 ^24^ we aimed for a minimum of 8 mice per group. Investigators were blinded to treatment group during quantification analysis. Results are expressed as mean ± standard error of the mean (SEM) and statistical analyses were performed (GraphPad Prism version 9.0.2; GraphPad, La Jolla, CA, USA) using Student’s t-test, or one-way or two-way ANOVA as detailed in figure legends. Multiple comparison analysis using Sidak’s post hoc tests were only performed when F<0.05 and there was no variance in homogeneity.

## Results

### EPA treatment resulted in accelerated reperfusion after HLI in mice

In olive oil-treated mice, the ischemic to non-ischemic flux ratio did not significantly improve from post-HLI values until day 10 and remained significantly lower than baseline (normal) until the final measurements at day 14. In comparison, across the treatment period, EPA treatment caused an accelerated reperfusion rate after HLI, where the recovery in the ischemic limb was faster, compared with mice treated with the olive oil (Figure 1A-C, p=0.018 treatment, p<0.0001 time). Mice treated with EPA had flux ratios higher than both post-HLI (p<0.001) and vehicle by day 7 (p=0.008) and the ischemic limb values were no longer different from baseline. In comparison, DHA had no effect on reperfusion when compared with vehicle-treated mice (Figure 1B,D, p=0.924 treatment, p<0.0001 time).

**Figure 1.**
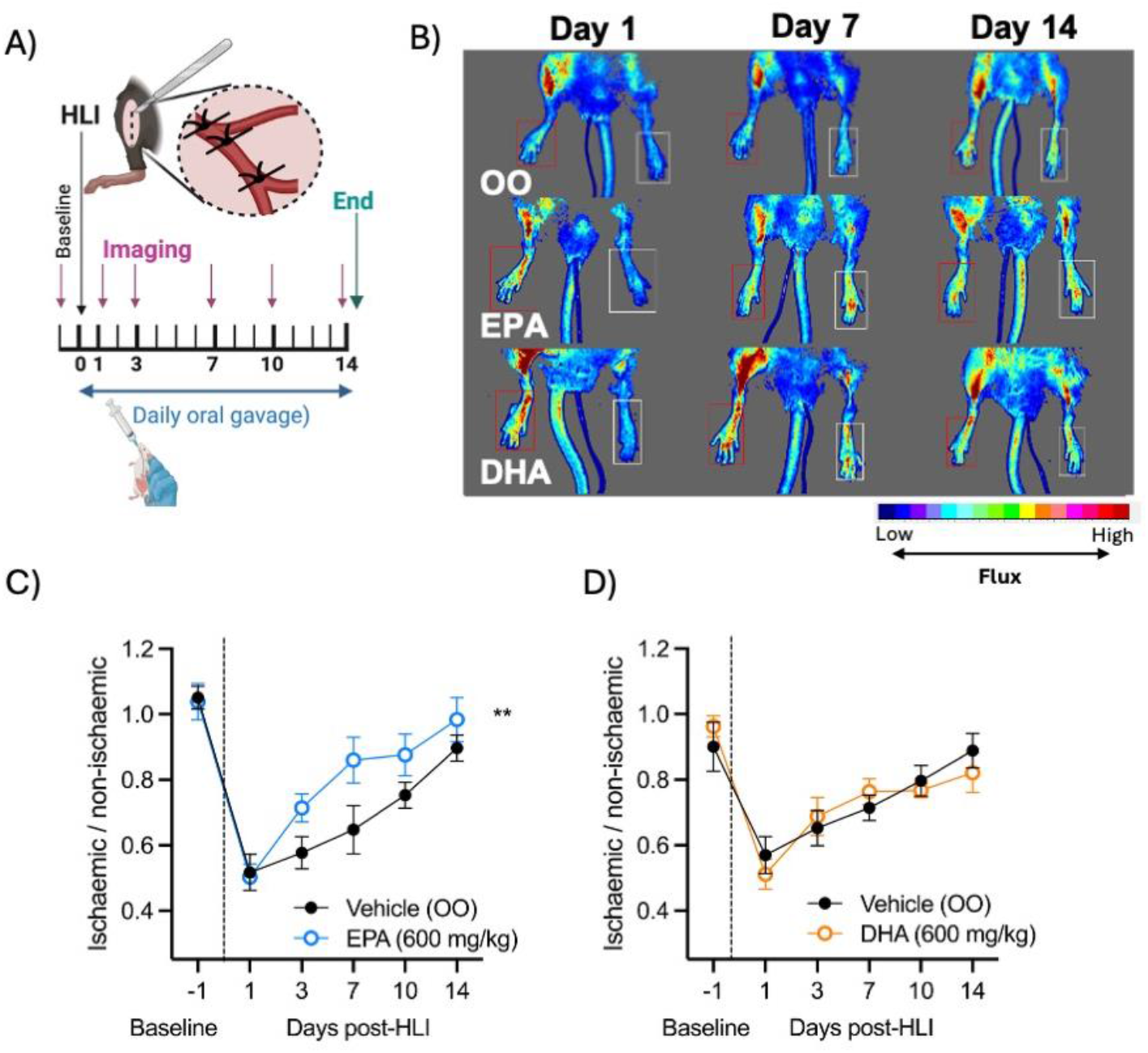
EPA promotes restoration of blood flow after hindlimb ischemia in C57BL/6 mice. **A)** Experimental timeline; **B)** Representative images showing Laser Doppler flux in hindlimbs from baseline (day -1) and post-hind limb ischemia (HLI) at days 1, 3, 7, 10 and 14 in mice treated with vehicle (olive oil, OO), EPA or DHA by daily oral gavage with 600 mg/kg of >99% pure omega 3 fatty acids, diluted in OO.) **C)** Summary data showing the reperfusion after HLI as ischaemic / non-ischaemic ratio in mice treated with vehicle (n=13) or EPA (n=13). **D)** Summary data showing the reperfusion after HLI as ischaemic / non-ischaemic ratio in mice treated with vehicle (n=8) or DHA (n=9). Data shown as mean ±SEM with statistical analysis by repeated measures two-way ANOVA with Sidak’s multiple comparison analysis. **P<0.01 vs. vehicle treatment.

### Angiogenesis is stimulated by EPA

Gastrocnemius muscle fibre morphology was assessed to measure compensatory muscle regeneration, indicative of angiogenesis ^26^. In all groups, HLI stimulated the centralisation of muscle fibre nuclei (Figure 2A-C). However compared with vehicle treated mice, EPA further enhanced regeneration of muscle fibres (Figure 2B), whereas this was not the case with DHA treatment (Figure 2C). Hindlimb fibrosis was quantified using picrosirius red staining and was found to be increased after HLI in mice treated with vehicle or DHA (Figure 2D-F). There was no effect of either EPA (Figure 2E) or DHA (Figure 2F) on fibrotic injury.

**Figure 2.**
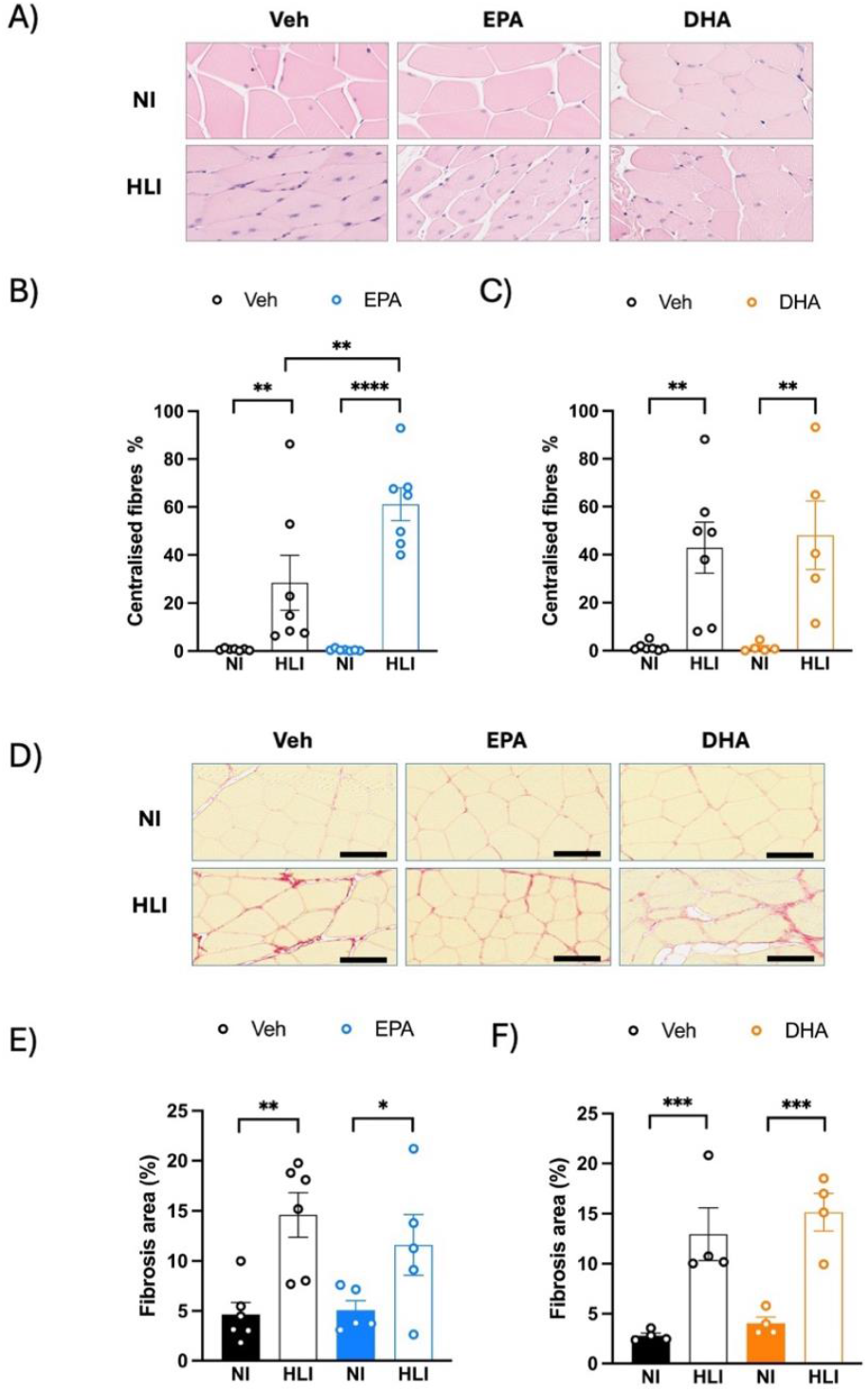
Muscle fibre regeneration is increased by EPA. **A)** H&E stain of cross-sections of gastrocnemius muscles for olive oil (OO)-, eicosapentaenoic acid (EPA)- and docosahexaenoic acid (DHA)-treated groups at 14 days post-ischemia, showing centralisation of nuclei (purple) within muscle fibres (pink) in tissue after hind limb ischemia (HLI). **B)** Muscle fibre regeneration was stimulated after HLI and further increased by EPA. **C)** DHA did not further increase muscle fibre regeneration. **D)** Representative images of gastrocnemius sections stained with picrosirius red showing increased collagen, indicative of muscle fibrotic injury occuring after HLI. **E)** EPA and DHA show similar amounts of fibrotic injury as vehicle controls. Data are presented as mean±SEM. Statistical analyses by two-way ANOVA with *P<0.05, **P<0.01, ***P<0.001, ****P<0.0001 determined by Sidak’s multiple comparison testing. Scale bars are equal to 50 µm.

To further support the role of EPA in promoting angiogenesis, gastrocnemius muscle sections were assessed by immunohistochemistry using a CD31 antibody, an established marker of endothelial cells and tissue vascularisation ^26^. By day 7, there was increased CD31 expression in the ischaemic hind limb of both vehicle and EPA-treated mice compared with their respective non-ischemic samples (Figure 3A, B). The increased post-ischemic vascularisation remained evident in both groups at day 14 (Figure 3C, D). EPA increased the number of CD31+ vessels after HLI, as compared to vehicle-treated mice at both day 7 and 14 (Figure 3B, D). The increased CD31 in EPA-treated HLI limbs was associated with greater eNOS protein expression in HLI muscle lysate from EPA-treated mice at day 14, an association not significant in the vehicle-treated mice (Figure 3 E, F). This demonstrates both eNOS upregulation by EPA after HLI, and increased vascularisation compared with vehicle treatment.

**Figure 3.**
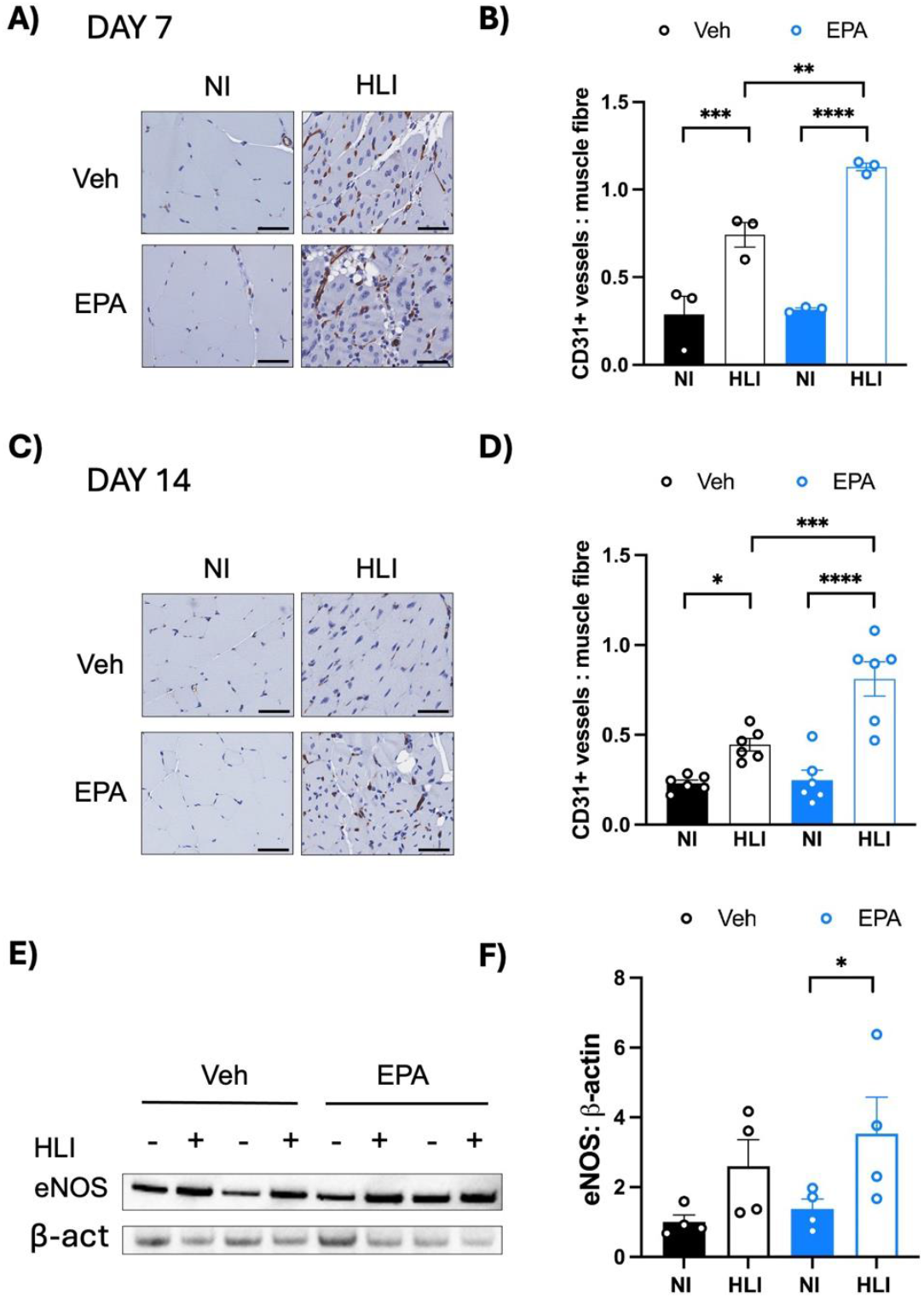
Neovascularisation in the ischemic limb increases after EPA treatment and is associated with increased eNOS protein expression. **A)** Representative images from day 7 showing vessels stained with CD31+ (brown) surrounding the muscle fibres. **B)** Quantification of the number of CD31+ stained vessels at day 7, shown as an average per mouse taken from multiple fields per section. **C)** Representative images from day 14 showing CD31+ vessels in brown. **D)** Quantification of the number of CD31+ stained vessels at day 14.**E)** Representative immunoblot of eNOS and β-actin **F)** Quantified eNOS expression relative to β-actin, expressed as a fold change of the Veh NI group. Data are presented as mean±SEM. Statistical analyses by two-way ANOVA with *P<0.05, **P<0.01, ***P<0.001, ****P<0.0001 determined by Sidak’s multiple comparison testing. Scale bars are equal to 50 µm.

### Hind limb HIF1α is increased early after HLI by EPA but other classical angiogenic pathways are not altered by EPA treatment

To determine whether expression of pro-angiogenic factors was altered, we performed quantitative real-time PCR to determine the expression of HIF1α and VEGF in gastrocnemius muscle homogenates. Temporal expression was determined by measuring samples collected from the mid-point (day 7) and endpoint (day 14) of the study. Compared with non-ischaemic tissue, HLI resulted in no change to HIF1α, but lower expression of VEGF at day 7 (Figure 4A, B). HIF1α was increased by day 14, whereas VEGF remained unchanged (Figure 4A, B. There was a significant relationship between omega 3 fatty acid treatment and HIF1α expression at day 7. While HIF1 α was not changed by DHA, there was a ∼40-fold increase in expression of HIF1α in EPA-treated mice, in both limbs (despite not reaching statistical significance in HLI, Figure 4A). This expression was then normalised by day 14 (Figure 4B), at the time that HIF1α increased in vehicle-treated mice. This early EPA-induced increase in HIF1α in the HLI limb, did not correlate with a change in VEGF (Figure 2B), and neither EPA nor DHA rescued the reduction in VEGF expression in the ischemic limb at day 7. VEGF expression in EPA-treated-mice was normal over the subsequent week, at the time when significant improvement in perfusion occurred in vehicle-treated mice. Taken together, there is a clear HIF1α response to omega-3 fatty acids but the extent of contribution to EPA-driven accelerated angiogenesis is unclear.

**Figure 4.**
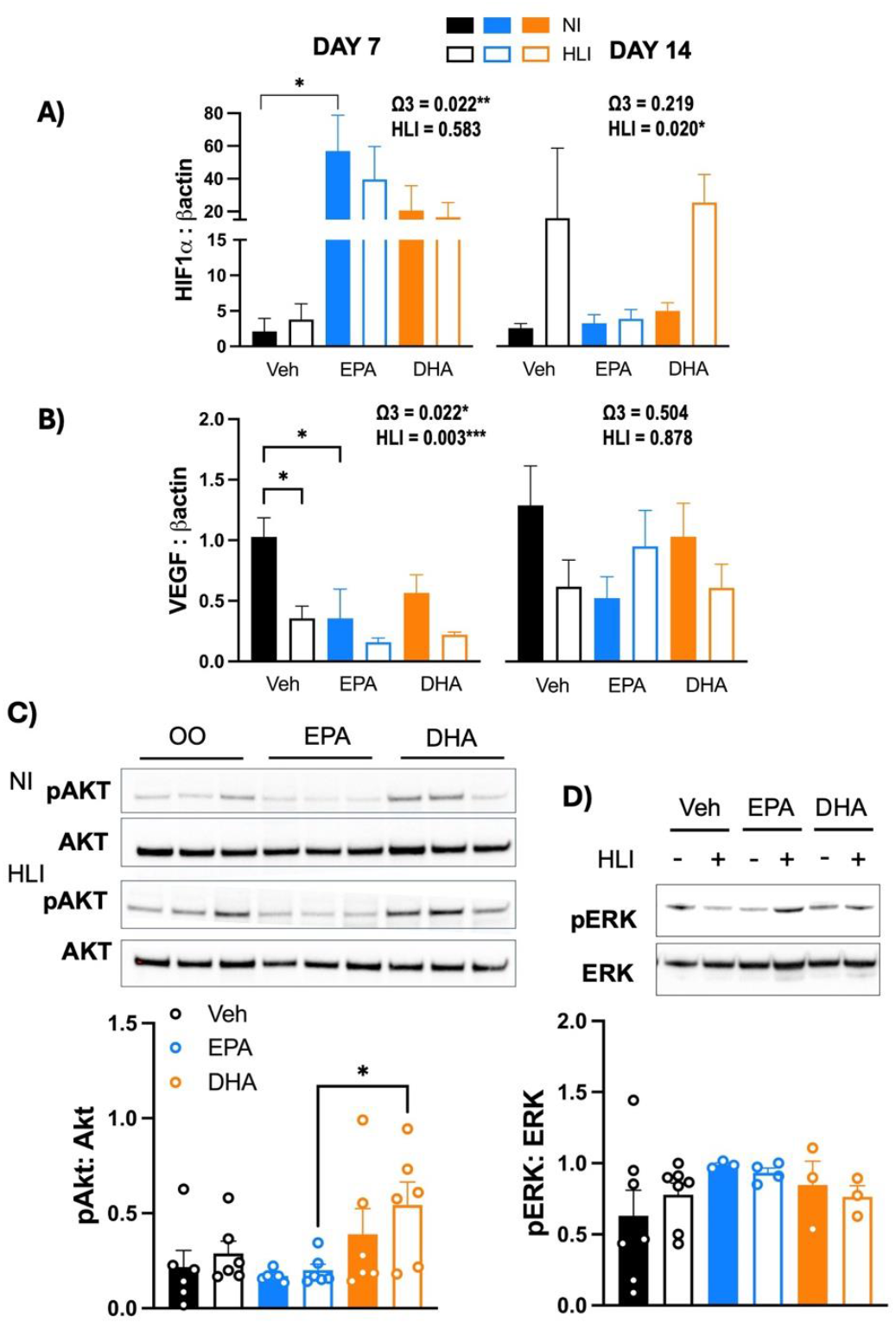
Common angiogenesis signaling pathways are not involved in EPA-mediated vascularisation. **A)** Hypoxia-inducible factor 1 α (HIF1α) and **B)** Vascular endothelial growth factor (VEGF) mRNA expression at day 7 and 14 in non-ischaemic (NI) vs. hind limb ischaemic (HLI) gastrocnemius muscle from mice treated with vehicle (olive oil, OO) vs. eicosapentaenoic acid (EPA, 600 mg/kg/day) or docosahexaenoic acid (DHA, 600 mg/kg/day). Data are normalised to β-actin, quantified relative to NI vehicle (fold change) and shown as mean ± SEM. n=3-4 at day 7 and n=6-7 at day 14. **C)** Phosphorylated AKT to AKT ratio and **D)** Phosphorylated ERK 1/2 to ERK 1/2 ratio in protein lysate of mice treated with vehicle, EPA or DHA. Statistical analysis by two-way analysis of variance to compare differences with omega 3 fatty acid treatment (Ω3) or after hindlimb ischemia (HLI). Sidak’s multiple comparison analysis tests were performed with *P<0.05.

We next quantified the activation of key pro-angiogenenic signaling molecules AKT and ERK by measuring phosphorylation using immunoblotting. There was no change in either of these pathways in HLI tissue at day 14 (Figure 4C-D). Interestingly, there was a relationship between omega-3 fatty acid treatment and phosphorylation of AKT (p=0.008) with DHA increasing the activation of AKT after HLI compared with EPA (p=0.025), although neither were significantly different from vehicle treatment. There were no changes in ERK phosphorylation with either EPA or DHA treatment. Taken together, these data suggest that neither signaling pathway is involved in EPA-mediated increases in reperfusion.

### EPA and DHA do not promote angiogenesis in human endothelial cells *in vitro*

To examine whether omega-3 fatty acids could promote angiogenesis in isolated human endothelial cells we used standard angiogenesis assays in HUVECs. The migration rate of HUVECs after scratch injury was similar in cells treated with vehicle (ethanol), EPA or DHA to controls (reduced serum media; Figure 5A-B), showing no effect of omega-3 fatty acids on cell migration. Likewise, tubule formation was not different in any treatment group, including a combination of EPA and DHA (Figure 5C-D). This suggests the beneficial effects of EPA on post-ischemic reperfusion are not attributable to simple endothelium stimulation but rather require more complex cellular interactions.

**Figure 5.**
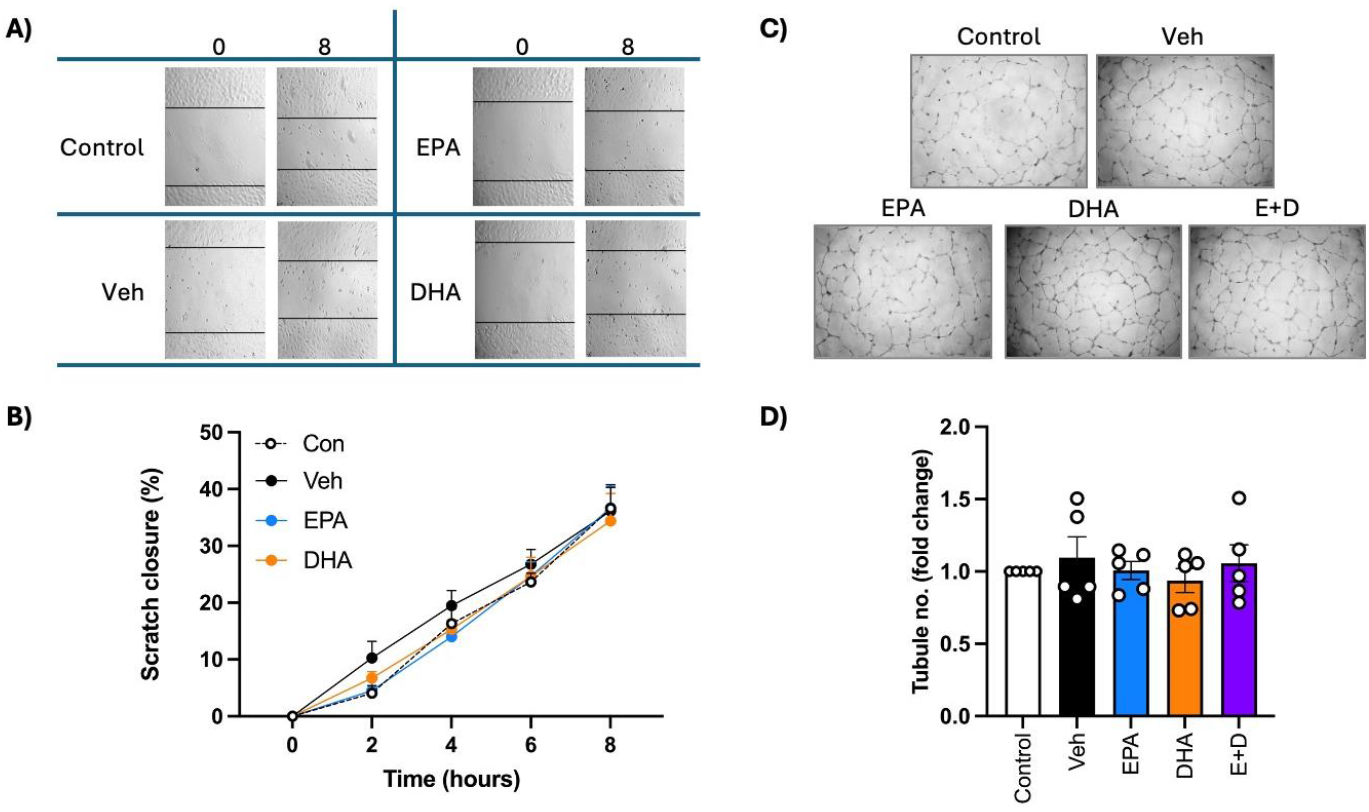
EPA and DHA do not promote angiogenesis in human umbilical vein endothelial cells in vitro. **A)** Representative images of human umbilical vein endothelial cells (HUVECs) showing migration after a scratch wound under control conditions (endothelial growth media (EGM2) diluted 1:3) or in response to treatment with vehicle (0.1% ethanol), EPA or DHA (0.01 mg/ml). **B)** Quantification of cell migration over 8 hours post-scratch, displaying no effect of EPA or DHA. **C)** Representative images of tubule formation in HUVECs grown on reduced growth factor extracellular matrix over 16 h in response to EGM2 (1:3), 0.1% ethanol, EPA or DHA (0.01 mg/ml) or combined EPA and DHA at 0.01 mg/ml each. Data are presented as mean±SEM. Statistical analyses by one-way (D) or two-way (B) ANOVA with Dunnett’s post-hoc analysis, n = 5 experiments with multiple technical replicates averaged for each experiment.

## Discussion

This study supports the hypothesis that EPA is superior to DHA in improving post-ischemic reperfusion. We demonstrated for the first time that high-dose EPA accelerates the restoration of blood flow to the ischemic limb, thereby supporting tissue regeneration through enhanced neovascularisation. Notably, this beneficial effect of EPA does not appear to be mediated through activation of classical angiogenic signaling pathways. In contrast, reperfusion, muscle regeneration and neovascularisation progressed at a similar rate in DHA- and vehicle-treated mice, indicating no additional benefit of DHA under these experimental conditions. Our work support and extend previous work demonstrating that pre-treatment with dietary omega-3 fatty acids, formulated with EPA and DHA at a 1.6:1 ratio, could enhance post-ischemic reperfusion ^23^.

To consider the translational potential of these findings, we recently undertook a systematic analysis of randomised controlled trials investigating omega-3 fatty acid supplementation in PAD ^11^. We concluded that there is currently insufficient evidence to support the use of omega-3 fatty acids in PAD or critical limb-threatening ischemia. However, significant limitations exist in the trials conducted to date, in agreement with a recent Cochrane review ^12^. These trials have not employed adequately high doses of omega-3 fatty acids (∼4 g/day) as recommended by the American Heart Association ^4^ and none have specifically focused on assessing functional improvement in PAD following high-dose EPA supplementation. Therefore, the pre-clinical findings of the current study provide further impetus for such trials to be conducted. In addition, a key strength of our study compared with previous work, is the use of a daily oral formulation of high-dose EPA and DHA. We calculated the 600 mg/kg/day dose for mice to be approximately equivalent to the recommended 4 g/day dose recommended in humans, a dosing strategy that has also been employed in prior studies from our group, and which demonstrated vascular anti-inflammatory effects^27^.

The mechanisms underpinning the effectiveness of EPA were explored. We identified increased eNOS protein at the study endpoint, which was associated with enhanced neovascularisation in ischemic hindlimbs of EPA-treated mice. A previous study identified improved eNOS bioavailability as a key mechanism by which an enriched omega-3 fatty acid diet confers vascular benefit, in agreement with our findings ^23^. We also demonstrated that omega-3 fatty acids increased HIF1α mRNA expression at the peak of accelerated reperfusion. However, although a ∼20-fold increase in HIF1α mRNA was observed, statistical significance in the context of low sample number was only reached in the non-ischaemic EPA-treated limb, suggesting a systemic effect of EPA. HIF1α stimulates the transcription of VEGF in response to hypoxia ^15^, yet we observed no change in VEGF mRNA expression with EPA treatment. This may be due to the timing of sample assessment relative to the rapid onset of the high-dose EPA administration, and it is possible that the peak mRNA response was missed. Given our observation of HIF1α and eNOS upregulation, it is likely that VEGF signalling was involved, and omega-3 fatty acids have been shown to stimulate hypoxia-induced VEGF expression ^20,23^. A mixed DHA and EPA formulation (1:1) has been shown to promote angiogenesis in placenta-derived mesenchymal stromal cells via upregulation of both fibroblast growth factor and VEGF-A, ^28^. The VEGF-promoting properties of omega-3 fatty acids have been demonstrated across a variety of cell types and vascular beds, yet there are also many studies reporting opposing effects of omega-3 fatty acids on angiogenesis. A key example is pathological retinal angiogenesis, a process key to vision impairment, which is inhibited by omega-3 fatty acids ^29-31^. In addition, there is substantial evidence showing inhibitory effects of omega-3 fatty acids on tumour angiogenesis ^18^. For example, omega-3 fatty acids suppress angiogenesis through reduced VEGF production in human colon cancer cells ^32^. Collectively, these findings suggest that omega-3 driven angiogenesis is an important mechanism for overcoming peripheral and cerebral ischemia, and the regional specificity of omega-3 fatty acid actions also provides neuroprotective and tumour anti-angiogenic effects, supporting the pharmaceutical application of omega-3 fatty acid formulations in a wide variety of clinical scenarios, particularly in an ageing population with multiple co-morbidities.

Whilst we demonstrated beneficial effects on EPA *in vivo*, we found no effect of EPA or DHA or their combination on angiogenesis, in isolated endothelial cells. This finding differs from a number of previously published studies. In HUVECs, conjugated EPA (generated via alkaline modification to induce additional conjugated double bonds) was shown to suppress VEGF-stimulated endothelial cell migration and sprouting angiogenesis, whereas pure EPA had no effect ^33^, consistent with our findings. In contrast, another study reported that *in vitro* treatment with EPA, but not DHA, inhibited VEGF-stimulated proliferation of isolated bovine carotid artery endothelial cells ^34^. Aside from angiogenesis, purified EPA has been shown to stimulate nitric oxide release from isolated porcine endothelial cells ^35^ and to improve eNOS bioavailability through interactions with lipid membrane caveolae ^36^. EPA can also protect HUVECs from inflammatory cytokine-dependent injury ^37^ and oxidised-LDL injury by improving eNOS bioavailability. The lack of effect observed in our *in vitro* studies may be explained by aspects of the experimental design including i) the absence of pre-treatment with EPA and ii) the lack of an injury stimulus such as inflammation or oxidative stress. With a modified experimental approach in future studies, it may be possible to expose a pro-angiogenic effect of EPA *in vitro*. An alternative explanation is that EPA requires a secondary mediator *in vivo* to promote angiogenesis.

We sought to compare the effects of EPA and DHA on common signaling pathways involved in promoting angiogenesis and therefore focused on assessing activation of the ERK1/2 and AKT pathways using phosphorylation-specific antibodies. We were surprised to find no effect of EPA on activation of either pathway, despite the significant stimulation of angiogenesis *in vivo*. Given that we also observed no effect of HLI on ERK or AKT activation, we acknowledge that the limitations related to the timing of sample collection and the inability to directly assess limb vascular endothelial cells, render our findings inconclusive. Future studies could address these limitations by incorporating additional tissue collection timepoints and performing cell-sorting approaches to isolate endothelial populations. EPA and DHA have been previously linked to AKT activation. A study in isolated endothelial progenitor cells showed that AKT mediated VEGF/eNOS-driven angiogenesis, an effect that was not observed with DHA ^38^. On the other hand, DHA has been shown to facilitate AKT nuclear translocation, conferring neuronal protection ^39^ and to activate AKT following cerebral ischemia ^40^. These findings are supported by our observation of greater AKT activation by DHA-treated ischemic limbs, although this was not associated with improved perfusion. The protective effects observed in retinopathy have also been largely attributed to DHA ^41^ Furthermore, DHA enhances the expression of eNOS and improves the bioavailability of nitric oxide in vascular endothelial cells ^42^. In addition, *apoE*^-/-^ mice supplemented with DHA exhibit reduced blood pressure and improved left ventricular function ^43^.

Another important consideration is that the vascular improvement observed with EPA, extended beyond angiogenesis alone. Omega-3 fatty acids have well-established antihypertensive effects ^44,45^ and are therefore understood to directly improve vascular function ^46^. A meta-analysis of 20 RCTs confirmed that purified EPA, but not DHA, significantly reduces systolic blood pressure. Flow-mediated dilatation, which is dependent on brachial artery endothelial function, can be improved by omega-3 fatty acids ^47^, including in patients with hypertension ^48^ and coronary artery disease ^49^, where endothelial function is impaired. Importantly, the latter study found improved flow mediated dilatation following treatment with a moderate dose of EPA (1.8g/day) over a 6-month period ^49^. Pre-clinical studies have probed these effects, commonly demonstrating vascular relaxation in both conduit and resistance arteries (reviewed in ^46^); an effect that is partially dependent on endothelial function, particularly through nitric oxide signalling ^50,51^, but also involves enhanced vascular smooth muscle relaxation driven by alternative mechanisms, such as ion channel regulation ^46,52^. Therefore, the improved blood flow observed with EPA in our study may have included a vasodilatory component; however, this cannot be easily assessed in the experimental model employed.

We have discussed above the constraints in elucidating the precise mechanisms by which EPA confers vascular benefit, due to limited sample collection timing and tissue specificity. Another key limitation of our study is that the HLI mouse model is predominately used to investigate angiogenesis and arteriogenesis ^53^ and there are inherent challenges in translating findings from this model to patients with PAD. A more clinically relevant approach would be to induce HLI in a complex metabolic model, for example using *apoE*^-/-^ mice subjected to a more severe ischemic insult via a two-stage procedure involving initial gradual femoral artery occlusion, followed by surgical excision of the femoral artery ^54^. Incorporating a diabetic model would also represent a relevant and important extension of the present study. Finally, inclusion of females in a follow up complex model is critical. Notably, the study by Turgeon and colleagues, showing improved post-ischemic reperfusion after an enriched fish oil diet, was conducted exclusively in female mice, whereas our study was performed in male mice. Taken together, these studies support a broader role for omega-3 fatty acids in ischemic vascular repair in both males and females.

In conclusion, we have identified the potential for high-dose EPA to be superior to DHA in promoting post-ischemic reperfusion in the hind limb. This effect appears to be driven, in part, by a direct role of EPA in enhancing neovascularisation, and further studies are warranted to elucidate the precise mechanisms underpinning its pro-angiogenic actions and to assess relevance in more complex disease models. These finding support the growing rationale for the use of purified EPA, rather than mixed omega-3 fatty acid formulations, in the clinical management of ischemic vascular diseases.

## Non-standard abbreviations

CVD: cardiovascular disease
DHA: docosahexaenoic acid
eNOS: endothelial nitric oxide synthase
EPA: eicosapentaenoic acid
ERK: extracellular signal-regulated kinase
HIF1α: hypoxia-inducible factor alpha
HLI: hind limb ischemia
HUVECs: Human umbilical vein endothelial cells
RCT: randomised controlled trial
VEGF: vascular endothelial growth factor

## Acknowledgements

The authors acknowledge the Monash Animal Research Platform for contributions to mouse supply and husbandry and the Monash Histology Platform for tissue sectioning and H&E and picrosirius red histological staining. Author contributions: Conceptualisation T.K.D, K.J,B, S,J,N; Study design T.K.D, K.J,B, S,J,N; Data collection T.K.D, K.J,B, A.P, G.B, J.T; Data analysis T.K.D, K.J,B, A.P, G.B, J.T; Interpretation of findings T.K.D, K.J,B, S,J,N; Drafting of manuscript T.K.D, K.J,B. Editing of final version all authors.

## References

1. Houghton JSM, Saratzis AN, Sayers RD, Haunton VJ. New Horizons in Peripheral Artery Disease. Age Ageing. 2024;53. doi: 10.1093/ageing/afae114

2. You Y, Wang Z, Yin Z, Bao Q, Lei S, Yu J, Xie X. Global disease burden and its attributable risk factors of peripheral arterial disease. Sci Rep. 2023;13:19898. doi: 10.1038/s41598-023-47028-5

3. Stone NJ. Fish consumption, fish oil, lipids, and coronary heart disease. Circulation. 1996;94:2337– 2340. doi: 10.1161/01.cir.94.9.2337

4. Skulas-Ray AC, Wilson PW, Harris WS, Brinton EA, Kris-Etherton PM, Richter CK, Jacobson TA, Engler MB, Miller M, Robinson JG. Omega-3 fatty acids for the management of hypertriglyceridemia: a science advisory from the American Heart Association. Circulation. 2019;140:e673–e691.

5. Bhatt DL, Steg PG, Miller M, Brinton EA, Jacobson TA, Ketchum SB, Doyle Jr RT, Juliano RA, Jiao L, Granowitz C. Cardiovascular risk reduction with icosapent ethyl for hypertriglyceridemia. New England Journal of Medicine. 2019;380:11–22.

6. Nicholls SJ, Lincoff AM, Garcia M, Bash D, Ballantyne CM, Barter PJ, Davidson MH, Kastelein JJ, Koenig W, McGuire DK. Effect of high-dose omega-3 fatty acids vs corn oil on major adverse cardiovascular events in patients at high cardiovascular risk: the STRENGTH randomized clinical trial. Jama. 2020;324:2268–2280.

7. Gans RO, Bilo HJ, Weersink EG, Rauwerda JA, Fonk T, Popp-Snijders C, Donker AJ. Fish oil supplementation in patients with stable claudication. The American journal of surgery. 1990;160:490–495.

8. Hammer A, Moertl D, Schlager O, Matschuck M, Seidinger D, Koppensteiner R, Steiner S. Effects of n-3 PUFA on endothelial function in patients with peripheral arterial disease: a randomised, placebo-controlled, double-blind trial. British journal of nutrition. 2019;122:698–706.

9. Carrero JJ, López-Huertas E, Salmerón LM, Ramos VE, Baró L, Ros E. Simvastatin and supplementation with ω-3 polyunsaturated fatty acids and vitamins improves claudication distance in a randomized PILOT study in patients with peripheral vascular disease. Nutrition research. 2006;26:637–643.

10. Ramírez-Tortosa C, López-Pedrosa JM, Suarez A, Ros E, Mataix J, Gil A. Olive oil-and fish oil-enriched diets modify plasma lipids and susceptibility of LDL to oxidative modification in free-living male patients with peripheral vascular disease: the Spanish Nutrition Study. British Journal of Nutrition. 1999;82:31–39.

11. Dao TK, Nerlekar N, Nicholls SJ, Bubb KJ. The effectiveness of intervention with omega-3 fatty acids, eicosapentaenoic and docosahexenoic acid in peripheral arterial disease: a systematic review and meta-analysis. Nutr Metab Cardiovasc Dis. 2025:104286. doi: 10.1016/j.numecd.2025.104286

12. Mohammady M, Brown T, Radmehr M, Shamsoddin E, Janani L. Omega-3 fatty acids for intermittent claudication. Cochrane Database Syst Rev. 2024;10:CD003833. doi: 10.1002/14651858.CD003833.pub5

13. Iyer SR, Annex BH. Therapeutic Angiogenesis for Peripheral Artery Disease: Lessons Learned in Translational Science. JACC Basic Transl Sci. 2017;2:503–512. doi: 10.1016/j.jacbts.2017.07.012

14. Hu B, Pei J, Wan C, Liu S, Xu Z, Zou Y, Li Z, Tang Z. Mechanisms of Postischemic Stroke Angiogenesis: A Multifaceted Approach. J Inflamm Res. 2024;17:4625–4646. doi: 10.2147/JIR.S461427

15. Forsythe JA, Jiang BH, Iyer NV, Agani F, Leung SW, Koos RD, Semenza GL. Activation of vascular endothelial growth factor gene transcription by hypoxia-inducible factor 1. Mol Cell Biol. 1996;16:4604– 4613. doi: 10.1128/MCB.16.9.4604

16. Shibuya M. Vascular Endothelial Growth Factor (VEGF) and Its Receptor (VEGFR) Signaling in Angiogenesis: A Crucial Target for Anti- and Pro-Angiogenic Therapies. Genes Cancer. 2011;2:1097–1105. doi: 10.1177/1947601911423031

17. Sakai C, Ishida M, Ohba H, Yamashita H, Uchida H, Yoshizumi M, Ishida T. Fish oil omega-3 polyunsaturated fatty acids attenuate oxidative stress-induced DNA damage in vascular endothelial cells. PloS one. 2017;12:e0187934.

18. Spencer L, Mann C, Metcalfe M, Webb MB, Pollard C, Spencer D, Berry D, Steward W, Dennison A. The effect of omega-3 FAs on tumour angiogenesis and their therapeutic potential. European journal of cancer. 2009;45:2077–2086.

19. Gonzalo-Gobernado R, Ayuso MI, Sansone L, Bernal-Jiménez JJ, Ramos-Herrero VD, Sánchez-García E, Ramos TL, Abia R, Muriana FJ, Bermúdez B. Neuroprotective effects of diets containing olive oil and DHA/EPA in a mouse model of cerebral ischemia. Nutrients. 2019;11:1109.

20. Wang J, Shi Y, Zhang L, Zhang F, Hu X, Zhang W, Leak RK, Gao Y, Chen L, Chen J. Omega-3 polyunsaturated fatty acids enhance cerebral angiogenesis and provide long-term protection after stroke. Neurobiol Dis. 2014;68:91–103. doi: 10.1016/j.nbd.2014.04.014

21. Lazzarin T, Martins D, Ballarin RS, Monte MG, Minicucci MF, Polegato BF, Zornoff L. The Role of Omega-3 in Attenuating Cardiac Remodeling and Heart Failure through the Oxidative Stress and Inflammation Pathways. Antioxidants (Basel). 2023;12. doi: 10.3390/antiox12122067

22. Heydari B, Abdullah S, Pottala JV, Shah R, Abbasi S, Mandry D, Francis SA, Lumish H, Ghoshhajra BB, Hoffmann U, et al. Effect of Omega-3 Acid Ethyl Esters on Left Ventricular Remodeling After Acute Myocardial Infarction: The OMEGA-REMODEL Randomized Clinical Trial. Circulation. 2016;134:378– 391. doi: 10.1161/CIRCULATIONAHA.115.019949

23. Turgeon J, Dussault S, Maingrette F, Groleau J, Haddad P, Perez G, Rivard A. Fish oil-enriched diet protects against ischemia by improving angiogenesis, endothelial progenitor cell function and postnatal neovascularization. Atherosclerosis. 2013;229:295–303.

24. Bubb KJ, Ravindran D, Cartland SP, Finemore M, Clayton ZE, Tsang M, Tang O, Kavurma MM, Patel S, Figtree GA. beta (3) Adrenergic Receptor Stimulation Promotes Reperfusion in Ischemic Limbs in a Murine Diabetic Model. Front Pharmacol. 2021;12:666334. doi: 10.3389/fphar.2021.666334

25. Thomas JM, Bamhare P, Mulangala J, Bursill CA, Nicholls SJ, Di Bartolo BA, Bubb KJ. Apolipoprotein C3 Promotes Angiogenesis in an Inflammatory Mouse Model of Peripheral Artery Disease. FASEB J. 2025;39:e71058. doi: 10.1096/fj.202502155R

26. Lee JJ, Arpino JM, Yin H, Nong Z, Szpakowski A, Hashi AA, Chevalier J, O’Neil C, Pickering JG. Systematic Interrogation of Angiogenesis in the Ischemic Mouse Hind Limb: Vulnerabilities and Quality Assurance. Arterioscler Thromb Vasc Biol. 2020;40:2454–2467. doi: 10.1161/ATVBAHA.120.315028

27. Pisaniello AD, Psaltis PJ, King PM, Liu G, Gibson RA, Tan JT, Duong M, Nguyen T, Bursill CA, Worthley MI. Omega-3 fatty acids ameliorate vascular inflammation: A rationale for their atheroprotective effects. Atherosclerosis. 2021;324:27–37.

28. Mathew SA, Bhonde RR. Omega-3 polyunsaturated fatty acids promote angiogenesis in placenta derived mesenchymal stromal cells. Pharmacological Research. 2018;132:90–98.

29. Connor KM, SanGiovanni JP, Lofqvist C, Aderman CM, Chen J, Higuchi A, Hong S, Pravda EA, Majchrzak S, Carper D, et al. Increased dietary intake of omega-3-polyunsaturated fatty acids reduces pathological retinal angiogenesis. Nat Med. 2007;13:868–873. doi: 10.1038/nm1591

30. Sapieha P, Stahl A, Chen J, Seaward MR, Willett KL, Krah NM, Dennison RJ, Connor KM, Aderman CM, Liclican E, et al. 5-Lipoxygenase metabolite 4-HDHA is a mediator of the antiangiogenic effect of omega-3 polyunsaturated fatty acids. Sci Transl Med. 2011;3:69ra12. doi: 10.1126/scitranslmed.3001571

31. Fu Z, Lofqvist CA, Shao Z, Sun Y, Joyal JS, Hurst CG, Cui RZ, Evans LP, Tian K, SanGiovanni JP, et al. Dietary omega-3 polyunsaturated fatty acids decrease retinal neovascularization by adipose-endoplasmic reticulum stress reduction to increase adiponectin. Am J Clin Nutr. 2015;101:879–888. doi: 10.3945/ajcn.114.099291

32. Calviello G, Di Nicuolo F, Gragnoli S, Piccioni E, Serini S, Maggiano N, Tringali G, Navarra P, Ranelletti FO, Palozza P. n-3 PUFAs reduce VEGF expression in human colon cancer cells modulating the COX-2/PGE 2 induced ERK-1 and-2 and HIF-1α induction pathway. Carcinogenesis. 2004;25:2303–2310.

33. Tsuzuki T, Shibata A, Kawakami Y, Nakagawa K, Miyazawa T. Conjugated eicosapentaenoic acid inhibits vascular endothelial growth factor-induced angiogenesis by suppressing the migration of human umbilical vein endothelial cells. The Journal of nutrition. 2007;137:641–646.

34. Yang SP, Morita I, Murota SI. Eicosapentaenoic acid attenuates vascular endothelial growth factor-induced proliferation via inhibiting Flk-1 receptor expression in bovine carotid artery endothelial cells. Journal of cellular physiology. 1998;176:342–349.

35. Boulanger C, Schini VB, Hendrickson H, Vanhoutte PM. Chronic exposure of cultured endothelial cells to eicosapentaenoic acid potentiates the release of endothelium-derived relaxing factor(s). Br J Pharmacol. 1990;99:176–180. doi: 10.1111/j.1476-5381.1990.tb14673.x

36. Li Q, Zhang Q, Wang M, Zhao S, Ma J, Luo N, Li N, Li Y, Xu G, Li J. Eicosapentaenoic acid modifies lipid composition in caveolae and induces translocation of endothelial nitric oxide synthase. Biochimie. 2007;89:169–177. doi: 10.1016/j.biochi.2006.10.009

37. Sherratt SCR, Libby P, Dawoud H, Bhatt DL, Mason RP. Eicosapentaenoic Acid Improves Endothelial Nitric Oxide Bioavailability Via Changes in Protein Expression During Inflammation. J Am Heart Assoc. 2024;13:e034076. doi: 10.1161/JAHA.123.034076

38. Chiu SC, Chiang EP, Tsai SY, Wang FY, Pai MH, Syu JN, Cheng CC, Rodriguez RL, Tang FY. Eicosapentaenoic acid induces neovasculogenesis in human endothelial progenitor cells by modulating c-kit protein and PI3-K/Akt/eNOS signaling pathways. J Nutr Biochem. 2014;25:934–945. doi: 10.1016/j.jnutbio.2014.04.007

39. Akbar M, Calderon F, Wen Z, Kim HY. Docosahexaenoic acid: a positive modulator of Akt signaling in neuronal survival. Proc Natl Acad Sci U S A. 2005;102:10858–10863. doi: 10.1073/pnas.0502903102

40. Eady TN, Belayev L, Khoutorova L, Atkins KD, Zhang C, Bazan NG. Docosahexaenoic acid signaling modulates cell survival in experimental ischemic stroke penumbra and initiates long-term repair in young and aged rats. PLoS One. 2012;7:e46151. doi: 10.1371/journal.pone.0046151

41. Fathima S, Prokopiou E, Georgiou T. Omega-3 Polyunsaturated Fatty Acids and Their Anti-Oxidant, Anti-Inflammatory and Neuroprotective Effects in Diabetic Retinopathy: A Narrative Review. Front Biosci (Landmark Ed). 2023;28:153. doi: 10.31083/j.fbl2807153

42. Balakumar P, Taneja G. Fish oil and vascular endothelial protection: bench to bedside. Free Radical Biology and Medicine. 2012;53:271–279.

43. Alfaidi MA, Chamberlain J, Rothman A, Crossman D, Villa-Uriol MC, Hadoke P, Wu J, Schenkel T, Evans PC, Francis SE. Dietary docosahexaenoic acid reduces oscillatory wall shear stress, atherosclerosis, and hypertension, most likely mediated via an IL-1–mediated mechanism. Journal of the American Heart Association. 2018;7:e008757.

44. Guo XF, Li KL, Li JM, Li D. Effects of EPA and DHA on blood pressure and inflammatory factors: a meta-analysis of randomized controlled trials. Crit Rev Food Sci Nutr. 2019;59:3380–3393. doi: 10.1080/10408398.2018.1492901

45. AbuMweis S, Jew S, Tayyem R, Agraib L. Eicosapentaenoic acid and docosahexaenoic acid containing supplements modulate risk factors for cardiovascular disease: a meta-analysis of randomised placebo-control human clinical trials. J Hum Nutr Diet. 2018;31:67–84. doi: 10.1111/jhn.12493

46. Bercea CI, Cottrell GS, Tamagnini F, McNeish AJ. Omega-3 polyunsaturated fatty acids and hypertension: a review of vasodilatory mechanisms of docosahexaenoic acid and eicosapentaenoic acid. British Journal of Pharmacology. 2021;178:860–877.

47. Wang Q, Liang X, Wang L, Lu X, Huang J, Cao J, Li H, Gu D. Effect of omega-3 fatty acids supplementation on endothelial function: a meta-analysis of randomized controlled trials. Atherosclerosis. 2012;221:536–543. doi: 10.1016/j.atherosclerosis.2012.01.006

48. Casanova MA, Medeiros F, Trindade M, Cohen C, Oigman W, Neves MF. Omega-3 fatty acids supplementation improves endothelial function and arterial stiffness in hypertensive patients with hypertriglyceridemia and high cardiovascular risk. J Am Soc Hypertens. 2017;11:10–19. doi: 10.1016/j.jash.2016.10.004

49. Sawada T, Tsubata H, Hashimoto N, Takabe M, Miyata T, Aoki K, Yamashita S, Oishi S, Osue T, Yokoi K, et al. Effects of 6-month eicosapentaenoic acid treatment on postprandial hyperglycemia, hyperlipidemia, insulin secretion ability, and concomitant endothelial dysfunction among newly-diagnosed impaired glucose metabolism patients with coronary artery disease. An open label, single blinded, prospective randomized controlled trial. Cardiovasc Diabetol. 2016;15:121. doi: 10.1186/s12933-016-0437-y

50. Omura M, Kobayashi S, Mizukami Y, Mogami K, Todoroki-Ikeda N, Miyake T, Matsuzaki M. Eicosapentaenoic acid (EPA) induces Ca(2+)-independent activation and translocation of endothelial nitric oxide synthase and endothelium-dependent vasorelaxation. FEBS Lett. 2001;487:361–366. doi: 10.1016/s0014-5793(00)02351-6

51. Zgheel F, Perrier S, Remila L, Houngue U, Mazzucotelli JP, Morel O, Auger C, Schini-Kerth VB. EPA:DHA 6:1 is a superior omega-3 PUFAs formulation attenuating platelets-induced contractile responses in porcine coronary and human internal mammary artery by targeting the serotonin pathway via an increased endothelial formation of nitric oxide. Eur J Pharmacol. 2019;853:41–48. doi: 10.1016/j.ejphar.2019.03.022

52. Limbu R, Cottrell GS, McNeish AJ. Characterisation of the vasodilation effects of DHA and EPA, n-3 PUFAs (fish oils), in rat aorta and mesenteric resistance arteries. PLoS One. 2018;13:e0192484– e0192484. doi: 10.1371/journal.pone.0192484

53. Yu J, Dardik A. A murine model of hind limb ischemia to study angiogenesis and arteriogenesis. Traumatic and Ischemic Injury: Methods and Protocols. 2018:135–143.

54. Krishna SM, Omer SM, Li J, Morton SK, Jose RJ, Golledge J. Development of a two-stage limb ischemia model to better simulate human peripheral artery disease. Scientific reports. 2020;10:3449.

